# Distinct neutralizing kinetics and magnitudes elicited by different SARS-CoV-2 variant spikes

**DOI:** 10.1101/2021.09.02.458740

**Authors:** Yang Liu, Jianying Liu, Jing Zou, Ping Ren, Scott C. Weaver, Xuping Xie, Pei-Yong Shi

## Abstract

The rapid evolution of SARS-CoV-2 mandates a better understanding of cross-protection between variants after vaccination or infection, but studies directly evaluating such cross-protection are lacking. Here we report that immunization with different variant spikes elicits distinct neutralizing kinetics and magnitudes against other SARS-CoV-2 variants. After immunizing hamsters with wild-type or mutant SARS-CoV-2 bearing variant spikes from Alpha, Beta, Gamma, or Epsilon, the animals developed faster and greater neutralization activities against homologous SARS-CoV-2 variants than heterologous variants, including Delta. The rank of neutralizing titers against different heterologous variants varied, depending on the immunized variant spikes. The differences in neutralizing titers between homologous and heterologous variants were as large as 62-, 15-, and 9.7-fold at days 14, 28, and 45 post-immunization, respectively. Nevertheless, all immunized hamsters were protected from challenges with all SARS-CoV-2 variants, including those exhibiting the lowest neutralizing antibody titers. The results provide insights into the COVID-19 vaccine booster strategies.

## Introduction

The global pandemic of severe acute respiratory syndrome coronavirus 2 (SARS-CoV-2) has caused >213 million infections and >4.4 million deaths (as of August 25, 2021 per https://coronavirus.jhu.edu/). Despite the unprecedented success of vaccine development for coronavirus disease 2019 (COVID-19),^1^ global control of the pandemic remains challenging because of insufficient vaccine production and vaccine hesitancy, as well as the emergence of new, more transmissible variants. Although coronaviruses have an intrinsic proofreading mechanism to maintain their long RNA genomes,^2^ SARS-CoV-2 continues to evolve, leading to the emergence of variants. Since viral spike protein is responsible for binding to the cellular receptor angiotensin-converting enzyme 2 (ACE2), SARS-CoV-2 variants have accumulated many of their mutations in the spike gene. Such spike mutations can alter transmission efficiency and/or immune escape. The first prevalent substitution that underwent a selective sweep, D614G, is located at the spike protein that enhances spike/ACE2 binding, making the virus more transmissible.^3–7^ Other substitutions, such as L452R and E484K in the spike receptor-binding domain (RBD), confer resistance of SARS-CoV-2 variants to therapeutic antibodies.^8,9^ Among the emerged variants, Beta (B.1.351) and Kappa (B.1.617.1) exhibit the least sensitivity to neutralization by immune sera from vaccinated people,^8,10–13^ whereas Alpha (B.1.1.7) and Delta (B.1.617.2) were associated with increased viral transmissibility.^14,15^ These observations have prompted the desire to modify the vaccine sequence to match variants of concern, such as Beta because of its reduced neutralization sensitivity to the current vaccine sera.^8,10^ However, one critical question about this modified vaccine approach is whether the new vaccine elicits potent neutralizing activities against other co-circulating variants. Along the same line, cross-protection among different variants after natural infection remains to be studied in unvaccinated populations.

## Results

To examine cross-protection among different variant spikes, we prepared a panel of four chimeric SARS-CoV-2 (**Extended Data Fig. 1a**), each bearing a distinct variant spike gene from Alpha (B.1.1.7), Beta (B.1.351), Gamma (P.1), or Epsilon (B.1.429) in the backbone of an early virus strain USA-WA1/2020 [isolated in January 2020 and defined as wild-type (WT)]. The four variants were selected based on their high prevalence at the onset of the project. Each variant spike contained a distinct set of mutations (**Fig. 1a**). An additional substitution E484K was added to the original Alpha variant (Alpha+E484K) as this mutation occurred in many clinical isolates.^16^ The spike genes from all recombinant viruses were sequenced to ensure no aberrant mutations. Comparable ratios of viral RNA copies versus plaque-forming units (RNA/PFU) were found for both WT and chimeric viruses when produced and analyzed on Vero E6 cells (**Extended Data Fig. 1b**), suggesting equivalent specific infectivity of the viral stocks.

**Figure 1.**
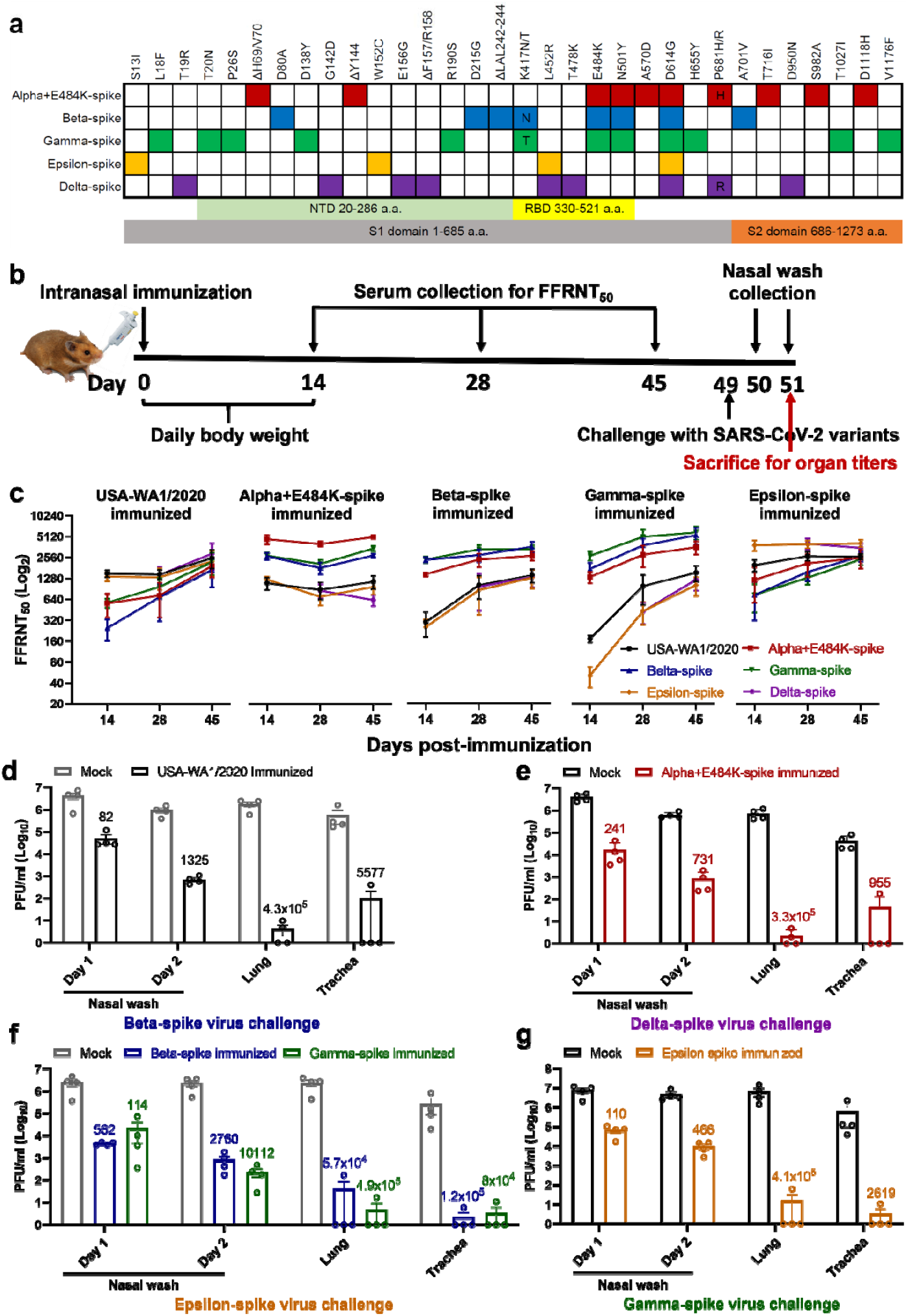
Variant spikes elicit neutralizing antibodies that cross-protect hamsters from challenges with SARS-CoV-2 variants. **a,** Amino acid substitutions in the spike protein among SARS-CoV-2 variants. The sequence of the spike from USA-WA1/2020 strain was used as a reference. NTD, N-terminal domain; RBD, Receptor binding domain. **b,** Experimental scheme of immunization and challenge in hamsters. The hamsters (n=4 per group) were intranasally immunized with 10^6^ PFU of WT or variant-spike SARS-CoV-2. Serum specimens were measured for FFRNT_50_ values on days 14, 28, and 45 post-immunization. On day 49 post-immunization, the hamsters were intranasally challenged by the indicated variant-spike SARS-CoV-2 (10^4^ PFU). The nasal washes were quantified for viral titers on days 1 and 2 post-challenge. All the hamsters were sacrificed on day 2 post-challenge for viral titers detection. **c,** Neutralization titers of hamster sera against SARS-CoV-2 spike variants on days 14, 28, and 45 post-immunization. Means ± standard errors of the mean are shown. **d-g,** Protection of immunized hamsters from the challenge of SARS-CoV-2 spike variants. The immunized hamsters and age-matched non-immunized hamsters (Mock) were challenged with selected variant viruses exhibiting the lowest neutralizing titers. The viral loads in the nasal wash (NW, days 1 and 2), lung, and trachea (day 2) were detected by plaque assays. The numbers above individual columns indicate the fold decrease in viral loads by comparing the means from the immunized group with that from the non-immunized mock group. Means ± standard errors of the mean are shown. The assay limit is 10 PFU/ml.

To analyze the immunogenicity of different variant spikes, we intranasally immunized hamsters with 10^6^ PFU of recombinant WT or variant-spike virus (**Fig. 1b**). The immunized animals developed different degrees of weight loss in the order of Alpha+E484K-spike > Beta-spike ≈ Gamma-spike > WT ≈ Epsilon-spike (**Extended Data Fig. 2a**). The weight loss results were consistent with the clinical scores, with the Alpha+E484K-spike virus causing the most severe disease (**Extended Data figure 2b**). These results suggest that Alpha+E484K-spike is the most pathogenic virus in the hamster model. Sera were collected on days 14, 28, and 45 post-immunization and measured for neutralizing titers against homologous and heterologous variant-spike viruses, including the currently prevalent Delta-spike virus (**Extended Data Fig. 1**). To increase assay throughput, we developed a “fluorescent foci” reduction neutralization test (FFRNT) by using mNeonGreen (mNG) reporter viruses (**Extended Data Fig. 3a**). The mNG gene was engineered into the open-reading-frame-7 (ORF7) of the viral genome.^17^ The protocols for the conventional plaque reduction neutralization test (PRNT) and FFRNT (**Extended Data Fig. 3b**) were similar except that the latter quantifies “fluorescent Foci” using a high-content imager in a high-throughput manner (**Extended Data Fig. 3c**). The two assays yielded comparable neutralizing titers for the same set of BNT162b2-vaccinated human sera (**Extended Data Fig. 3d,e**), validating the utility of FFRNT for neutralization test.

FFRNT analysis of immunized hamster sera showed distinct neutralizing profiles against homologous and heterologous SARS-CoV-2 variants (Summary in **Fig. 1c** and details in **Extended Data Fig. 4** and **Extended Data Tables 1-3**). (i) Each variant spike elicited faster and higher neutralizing titers against its homologous SARS-CoV-2 variant than heterologous variants; (ii) The magnitudes and ranks of neutralizing titers against different heterologous variants varied depending on the immunized variant spikes; (iii) Unlike other variant spike-immunized groups, the Alpha-spike-immunized animals did not seem to increase the neutralizing titers against heterologous variants from days 14 to 45 post-immunization. It is notable that from days 14 to 45 post-immunization, homologous neutralization titers increased by ≤ 2.32-fold, whereas heterologous neutralization titers could increase up to 22-fold when Gamma-spike-immunized sera were tested against epsilon-spike SARS-CoV-2 (**Fig. 1c**). On days 14, 28, and 45 post-immunization, the differences in neutralizing titers between homologous and heterologous variants could be as large as 62-, 15-, 9.7-fold, respectively (**Fig. 1c**). Collectively, the results demonstrate that vaccination of hamsters with different variant spikes elicits distinct kinetics, magnitudes, and ranks of neutralizing titers against homologous and heterologous SARS-CoV-2 variants.

To directly evaluate cross-protection, we selected variant viruses exhibiting the lowest neutralizing titers for each immunized group to challenge the hamsters on day 49 post-immunization. Specifically, animals immunized with WT, Alpha-, Beta-, Gamma-, or Epsilon-spike were challenged with 10^4^ PFU of Beta-, Delta-, Epsilon-, Epsilon-, and Gamma-spike SARS-CoV-2, respectively. Compared with PBS-immunized, challenged animals, all variant spike-immunized hamsters were protected from the challenge and developed significantly lower viral loads in nasal washes (82- to 10,112-fold), tracheas (955- to 120,000-fold), and lungs (57,000- to 490,000-fold) (**Fig. 1d**).

## Discussion

Our study provided experimental evidence against the need to modify vaccine sequences to match the currently circulating SARS-CoV-2 variants of concern. Our results showed distinct cross-neutralizing profiles elicited by different variant spikes, underscoring the heterogenicity in neutralization titers against different variants for any modified spike vaccines. Such modified vaccines may pose logistic challenges for vaccine implementation because (i) multiple variants often cocirculate and (ii) the constellation of variants may differ at different geographic regions and change rapidly over time, such as the recent replacement of the Alpha by the Delta variant in many regions; these replacements have generally not been predictable. Although different vaccine choices should be prescribed depending on the prevalence of specific variants, the prescribed vaccine should also be effective against other co-circulating variants. Our results, together with the observation that BNT612b2-immunized sera remained active in neutralizing all tested variants,^10–12^ support the strategy to continue the currently approved BNT612b2 vaccine for global immunization. This strategy is further bolstered by the real-world effectiveness of two BNT612b2 doses at efficacy rates of 89.5%, 75%, and 88% against Alpha, Beta, and Delta variants, respectively^18,19^. When protective immunity wanes over time, a third BNT612b2 booster could be administered to enhance the overall neutralizing titers to prevent infection and disease due to new variants. However, this strategy is contingent on the sensitivity of future variants to the immunity elicited by the current vaccine. As herd immunity continues to increase through natural infection and vaccination, selective pressures for the evasion of immunity may rise. The long-term strategy should include (i) surveillance of immune escape of new variants and (ii) preparedness for changes to vaccine strains with immune escape capability.

A limitation of this study is the use of chimeric viruses rather than the use of clinically approved vaccine platforms for expressing variant spikes or clinical variant isolates for the challenge. The neutralizing profile elicited by chimeric viruses may differ from that elicited by the clinically approved vaccine platforms. In chimeric virus-immunized hamsters, immune responses to non-spike viral proteins may provide added protection when compared with animals immunized with spike-alone vaccines such as mRNA and adenovirus-expression platforms. Despite this limitation, it is conceivable that the relative rank of neutralizing levels would be preserved against different SARS-CoV-2 variants.

In summary, increasing global immunization with the currently available safe and effective vaccines, together with boosters when needed, is the strategy to end the COVID-19 pandemic. The design of the booster vaccines depends on whether the newly emerged variants can escape the immunity generated by the current vaccines or natural infections. Potential immune escape of any new variants should be closely monitored by laboratory studies and real-world breakthroughs in vaccinated and infected individuals.

## Methods

### Ethics statement

Hamster studies were performed under the guidance of the Care and Use of Laboratory Animals of the University of Texas Medical Branch (UTMB). The protocol was approved by the Institutional Animal Care and Use Committee (IACUC) at UTMB. All the hamster operations were performed under anesthesia by isoflurane to minimize animal suffering.

### Animals and Cells

The Syrian golden hamsters (HsdHan:AURA strain) were purchased from Envigo (Indianapolis, IN). Vero E6 cells, an African green monkey kidney epithelial cell line (ATCC, Manassas, VA, USA), were cultured in Dulbecco’s modified Eagle’s medium (DMEM; Gibco/Thermo Fisher, Waltham, MA, USA) with 10% fetal bovine serum (FBS; HyClone Laboratories, South Logan, UT) plus 1% ampicillin/streptomycin (Gibco). The authenticity of Vero E6 cells was verified using Short Tandem Repeat profiling by ATCC. The cells were tested negative for mycoplasma.

### Construction of chimeric SARS-CoV-2s with variant spikes and mNeonGreen (mNG) reporter viruses

All spike mutations from different variants were engineered into an infectious cDNA clone of an early SARS-CoV-2 isolate USA-WA1/2020 using a standard PCR-based mutagenesis method. The protocol for the construction of recombinant SARS-CoV-2 was reported previously.^17,20^ To construct the mNG reporter viruses with variant spikes, the mNG gene was engineered into the open-reading-frame-7 (ORF7) of the viral genome. The full-length cDNAs of the viral genome containing the variant spike mutations were assembled by in vitro ligation. The resulting genome-length cDNAs served as templates for in vitro transcription of full-length viral RNAs. The full-length viral RNA transcripts were electroporated into Vero E6 cells. On day 2 post electroporation (when the electroporated cells developed cytopathic effects due to recombinant virus production and replication), the original viral stocks (P0) were harvested from the culture medium. The P0 viruses were amplified on Vero E6 cells for another round to produce working viral stocks (P1). The complete spike genes from the P1 viruses were sequenced to ensure no undesired mutations. The P1 viruses were used for the following study.

### Plaque assay

Approximately 1.2×10^6^ Vero E6 cells were seeded to each well of 6-well plates and cultured at 37°C, 5% CO_2_ for 16 h. The virus was serially diluted in DMEM with 2% FBS and 200 μl diluted viruses were transferred onto the monolayer of Vero E6 cells. The viruses were incubated with the cells at 37°C with 5% CO_2_ for 1 h. After the incubation, 2 ml of overlay medium (DMEM medium supplemented with 1% agar) was added to the infected cells per well. The overlay medium contained DMEM with 2% FBS, 1% penicillin/streptomycin, and 1% sea-plaque agarose (Lonza, Walkersville, MD). After a 2-day incubation, plates were stained with neutral red (Sigma-Aldrich, St. Louis, MO) and plaques were counted on a lightbox.

### Quantitative real-time RT-PCR assays

RNA copies of SARS-CoV-2 samples were detected by quantitative real-time RT-PCR (RT-qPCR) assays were performed using the iTaq SYBR Green One-Step Kit (Bio-Rad) on the LightCycler 480 system (Roche, Indianapolis, IN) following the manufacturer’s protocols. The absolute quantification of viral RNA was determined by a standard curve method using an RNA standard (*in vitro* transcribed 3,480 bp containing genomic nucleotide positions 26,044 to 29,883 of SARS-CoV-2 genome).

### Hamster infections

Four- to six-week-old male golden Syrian hamsters, strain HsdHan:AURA (Envigo, Indianapolis, IN), were intranasally immunized with 10^6^ PFU recombinant WT or variant spike virus on day 0. The immunized animals were weighed and monitored for signs of illness daily. Sera were collected on days 14, 28, and 45 post-immunization and measured for neutralizing titers against homologous and heterologous variant-spike viruses. On day 49, animals from each immunized group were challenged with 10^4^ PFU of selected variant viruses exhibiting the lowest neutralizing titers. Specifically, animals immunized with WT, Alpha-, Beta-, Gamma-, or Epsilon-spike were challenged with the Beta-, Delta-, Epsilon-, Epsilon-, and Gamma-spike SARS-CoV-2, respectively. Nasal washes were collected in 400 μl sterile DPBS at indicated time points. Animals were humanely euthanized for organ collections after 2 days of the challenge. The harvested tracheae and lungs were placed in a 2-ml homogenizer tube containing 1 ml of maintenance media (DMEM supplemented with 2% FBS and 1% penicillin/streptomycin) and stored at −80°C. Samples were subsequently thawed, lung or tracheae were homogenized using TissueLyser II (Qiagen, Hilden, Germany) for 1 min at 26 sec-1, and debris was pelleted by centrifugation for 5 min at 16,100×g. Infectious titers were determined by plaque assay.

### Human serum specimens

The research protocol regarding the use of human serum specimens was reviewed and approved by the University of Texas Medical Branch (UTMB) Institutional Review Board. The approved IRB protocol number is 20-0070. All human serum specimens were obtained from the vaccinated subjects at the UTMB. All specimens were de-identified from patient information.

### Fluorescent foci reduction neutralization assay

Neutralization titers of human and hamster sera were measured by fluorescent foci reduction neutralization assay (FFRNT) using the mNG SARS-CoV-2. Briefly, Vero E6 cells (2.5 × 10^4^) were seeded in each well of black CLEAR flat-bottom 96-well plate (Greiner Bio-one™). The cells were incubated overnight at 37°C with 5% CO_2_. On the following day, each serum was 2-fold serially diluted in the culture medium with the first dilution of 1:10. The diluted serum was incubated with 100 PFU of mNG SARS-CoV-2 at 37 °C for 1 h (final dilution range of 1:20 to 1:5120), after which the serum-virus mixtures were inoculated onto Vero E6 cell monolayer in 96-well plates. After 1 h of infection, the inoculum was removed and 100 μl of overlay medium (DMEM supplemented with 0.8% methylcellulose, 2% FBS, and 1% P/S) was added to each well. The plates were incubated at 37°C for 20 h. The raw images were acquired using Cytation™ 7 (BioTek) armed with 2.5× objective and processed using the default software setting. The foci in each well were counted and normalized to the non-serum-treated controls to calculate the relative infectivities. The curves of the relative infectivity versus the serum dilutions (log10 values) were plotted using Prism 9 (GraphPad). A nonlinear regression method was used to determine the dilution fold that neutralized 50% of mNG SARS-CoV-2 (defined as FFRNT). Each serum was tested in duplicates.

### Plaque reduction neutralization test (PRNT)

A conventional 50% plaque-reduction neutralization test (PRNT_50_) was performed to measure the serum-mediated virus suppression as reported previously^21^. Individual sera were 2-fold serially diluted in culture medium with a starting dilution of 1:40 (dilution range of 1:40 to 1:1280). The diluted sera were incubated with 100 PFU of USA-WA1/2020 (WT) or mutant SARS-CoV-2. After 1 h incubation at 37°C, the serum-virus mixtures were inoculated onto 6-well plates with a monolayer of Vero E6 cells pre-seeded on the previous day. The minimal serum dilution that suppressed >50% of viral plaques is defined as PRNT_50_.

## Data availability

The data that support the findings of this study are available from the corresponding authors upon reasonable request.

## Acknowledgments

We thank Phillip R. Dormitzer for his helpful discussions during the study. P.-Y.S. was supported by NIH grants HHSN272201600013C, AI134907, AI145617, and UL1TR001439, and awards from the Sealy & Smith Foundation, the Kleberg Foundation, the John S. Dunn Foundation, the Amon G. Carter Foundation, the Gilson Longenbaugh Foundation, and the Summerfield Robert Foundation. S.C.W. was supported by NIH grant R24 AI120942. P.R. and X.X. were partially supported by the Sealy & Smith Foundation. J.L. was supported by James W. McLaughlin Fellowship Fund.

## Author contributions

Conceptualization, Y.L., S.C.W., X.X., P.-Y.S.; Methodology, Y.L. J.L., J.Z., X.X., P.-Y.S; Investigation, Y.L., J.L., J.Z., S.C.W., X.X., P.-Y.S.; Resources, P.R., S.C.W., P.-Y.S.; Data Curation, Y.L., J.L., J.Z., X.X., P.-Y.S.; Writing-Original Draft, Y.L., X.X., P.-Y.S.; Writing-Review & Editing, Y.L., J.L., J.Z., P.R., S.C.W., X.X., P.-Y.S.; Supervision, S.C.W., X.X., P.-Y.S.; Funding Acquisition P.-Y.S..

---

**Extended Data Figure 1.**
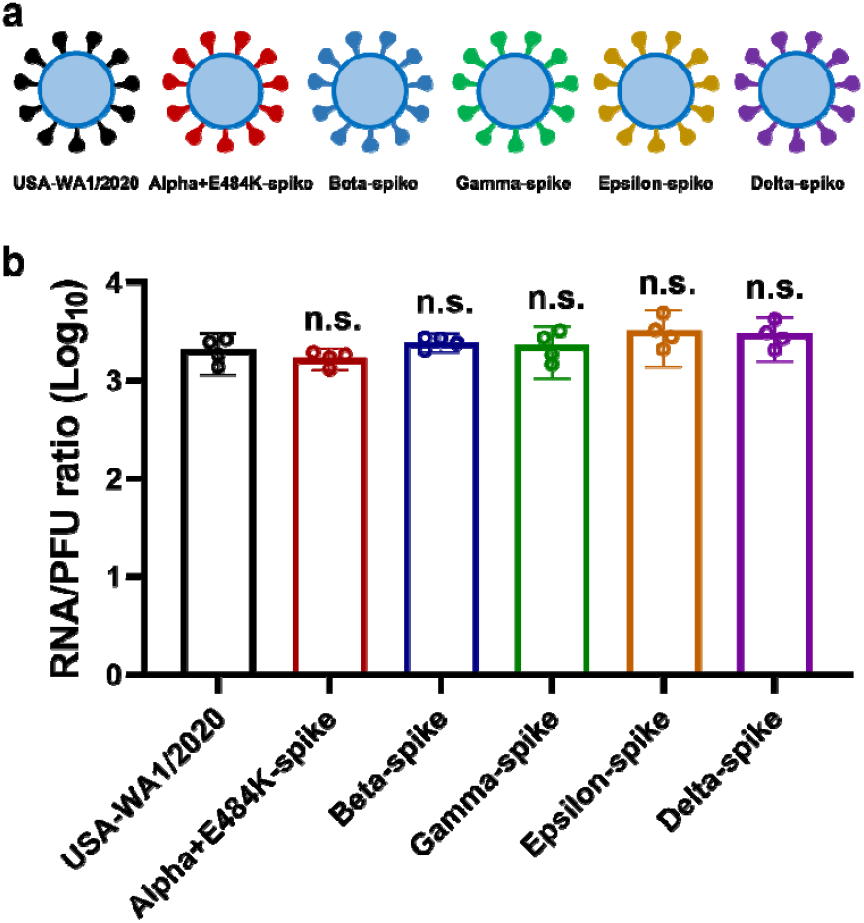
The RNA/PFU ratios of different SARS-CoV-2 variants. **a,** Diagram of SARS-CoV-2 spike variants. The spike genes from Alpha, Beta, Gamma, Epsilon, and Delta variants of SARS-CoV-2 were introduced into USA-WA1/2020 backbone. **b,** Ratios of viral genomic RNA versus plaque-forming unit (RNA/PFU) of SARS-CoV-2 spike variants. The genomic RNA and PFU of individual viral stocks were measured by RT-qPCR and plaque assay, respectively. The USA-WA1/2020 strain served as a control. Dots represent individual biological replicates from 4 aliquots of viruses. The means with 95% confidence intervals are shown. A non-parametric Mann-Whitney test was used to determine significant differences between USA-WA1/2020 and other variants. *P* values were adjusted using the Bonferroni correction to account for multiple comparisons. Differences were considered significant if *P* < 0.05; n.s., no statistical difference.

**Extended Data Figure 2.**
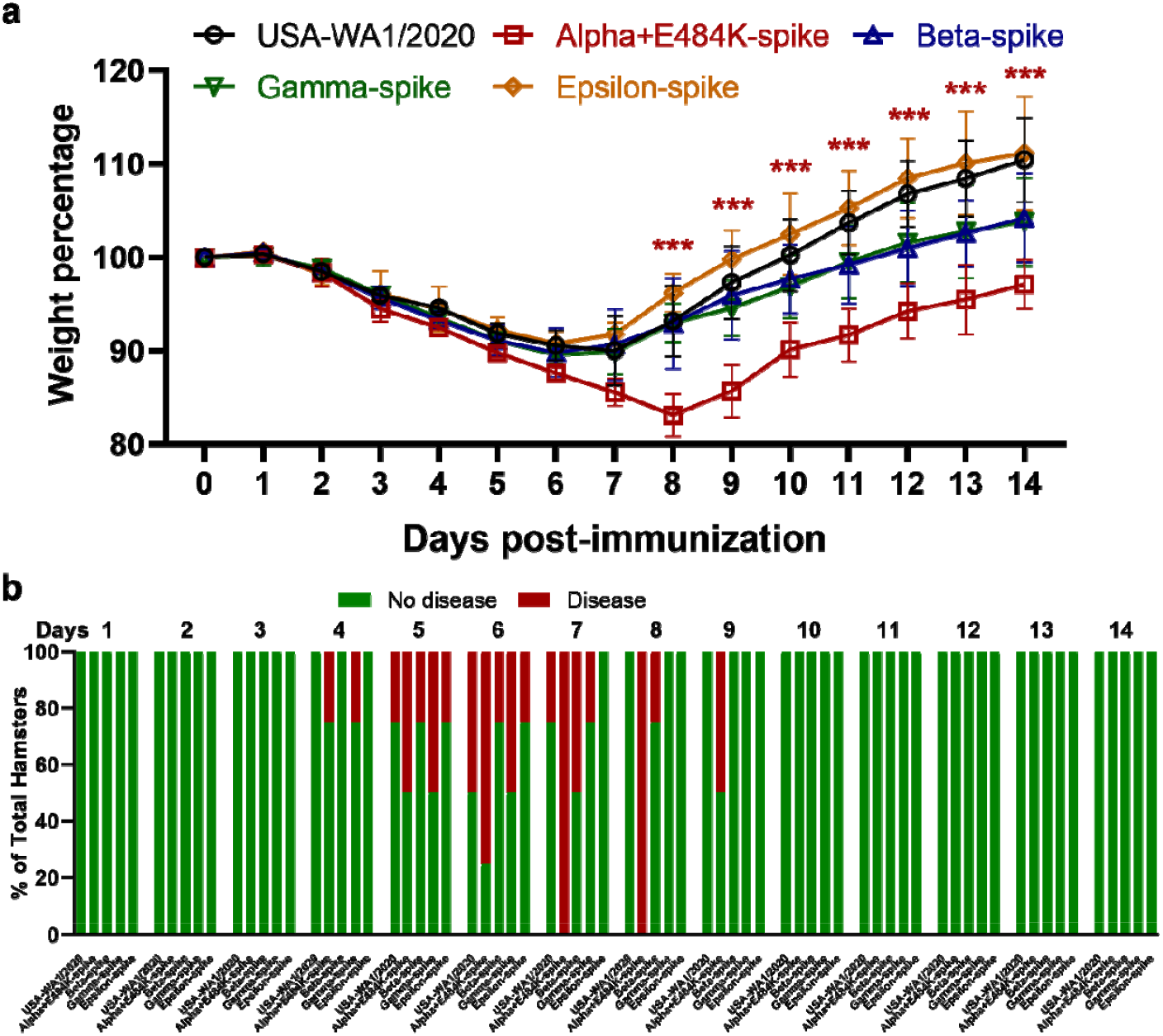
Morbidity of hamsters after immunized with variant-spike SARS-CoV-2. **a,** Hamster body weight loss after immunized with variant-spike SARS-CoV-2. The hamsters (n=4) were intranasally infected with 10^6^ PFU viruses. The body weights were measured daily from day 0 to day 14 days post-immunization. The weight loss data are shown as mean ± standard deviation and statistically analyzed using two-way ANOVA Turkey’s multiple comparisons. The red stars show the statistical significance (*** *P* < 0.001) between USA-WA1/2020-immunized hamsters and Alpha+E484K-spike-immunized hamsters. **b,** Percentages of hamsters with or without diseases (including ruffled fur, lethargic, hunched, and reluctance to move when stimulated) from day 1 to day 14 post-immunization.

**Extended Data Figure 3.**
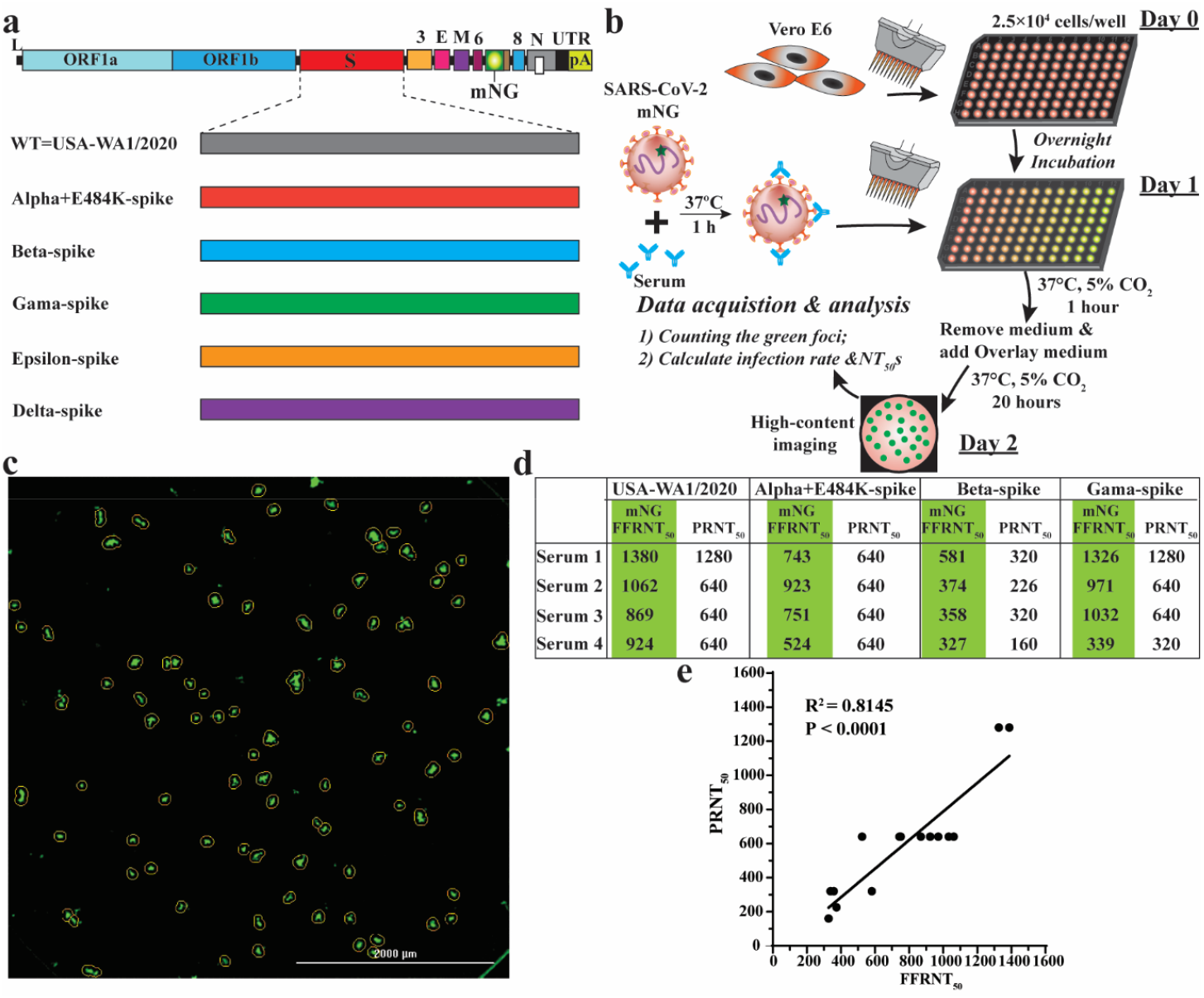
Correlation between FFRNT_50_ and PRNT_50_. **a,** Diagram of mNG USA-WA1/2020 and spike variants. mNG, mNeongreen fluorescence protein gene. **b,** Workflow of fluorescent foci reduction neutralization (FFRNT) assay. The details of the FFRNT assay were described in the Methods. **c,** Representative images of foci formed in a 96-well plate after 20 h of infection. **d,** FFRNT_50_, and PRNT_50_ values for four human sera. The FFRNT_50_ values are shaded in green. **e,** Correlation of FFRNT_50_ and PRNT_50_. The Pearson’s correlation coefficients and *P* values (two-tailed) are indicated.

**Extended Data Figure 4.**
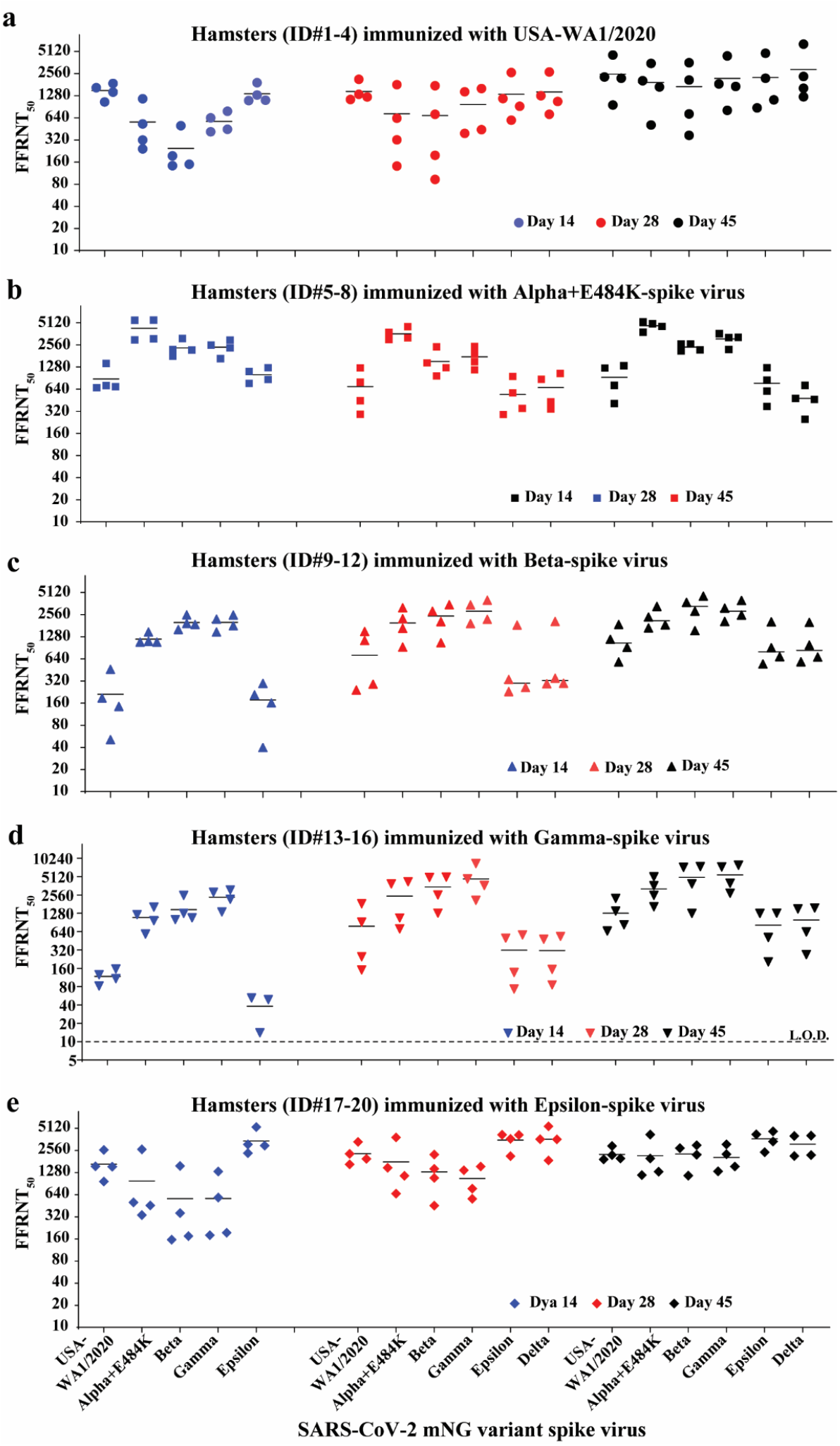
FFRNT_50_s of hamster sera against mNG SARS-CoV-2 spike variants on days 14, 28, and 45 post-immunization. **a-e,** Hamster (n=4 per group) were immunized with WT USA-WA1/2020 (**a**), Alpha+E484K-spike virus (**b**), Beta-spike virus (**c**), Gamma-spike virus (**d**), Epsilon-spike virus (**e**). Sera were collected on days 14, 28, and 45 post-immunization and tested for neutralizing activities against the indicated mNG viruses by FFRNT. The original FFRNT_50_ values are presented in **Extended Data Tables 1-3.**

**Extended Data Table 1.** FFRNT_50_s of twenty hamster sera on day 14 post-immunization.

| Virus for infection | Hamster ID | FFRNT<sub>50</sub>s against SARS-CoV-2 spike variants |  |  |  |  |
| --- | --- | --- | --- | --- | --- | --- |
|  |  | USA-WA1/2020 | Alpha+E484K-spike | Beta-spike | Gamma-spike | Epsilon-spike |
|  |  | mNG | mNG | mNG | mNG | mNG |
| USA-WA1/2020 | 1 | 1048 | 320 | 194 | 446 | 1113 |
|  | 2 | 1875 | 1158 | 498 | 785 | 1916 |
|  | 3 | 1643 | 242 | 143 | 413 | 1304 |
|  | 4 | 1426 | 528 | 149 | 637 | 1099 |
|  | Mean | 1498 | 562 | 246 | 570 | 1358 |
| Alpha+E484K-spike | 5 | 689 | 5561 | 1789 | 1660 | 1111 |
|  | 6 | 669 | 5526 | 2214 | 2543 | 864 |
|  | 7 | 1431 | 2981 | 3122 | 2967 | 762 |
|  | 8 | 714 | 3087 | 2175 | 2318 | 1246 |
|  | Mean | 876 | 4289 | 2325 | 2372 | 996 |
| Beta-spike | 9 | 461 | 1079 | 1610 | 2226 | 296 |
|  | 10 | 188 | 1482 | 2537 | 2520 | 161 |
|  | 11 | 51 | 1082 | 1947 | 1486 | 40 |
|  | 12 | 144 | 1113 | 1881 | 1802 | 208 |
|  | Mean | 211 | 1189 | 1994 | 2009 | 176 |
| Gamma-spike | 13 | 108 | 1215 | 2525 | 2833 | 49 |
|  | 14 | 125 | 583 | 1006 | 1346 | <10 |
|  | 15 | 155 | 967 | 1075 | 2166 | 14 |
|  | 16 | 82 | 1617 | 1275 | 3092 | 52 |
|  | Mean | 118 | 1096 | 1470 | 2359 | 38 |
| Epsilon-spike | 17 | 966 | 337 | 156 | 194 | 2346 |
|  | 18 | 2602 | 502 | 358 | 583 | 5297 |
|  | 19 | 1535 | 457 | 175 | 179 | 2977 |
|  | 20 | 1552 | 2637 | 1572 | 1321 | 3085 |
|  | Mean | 1664 | 983 | 565 | 569 | 3426 |

**Extended Data Table 2.**
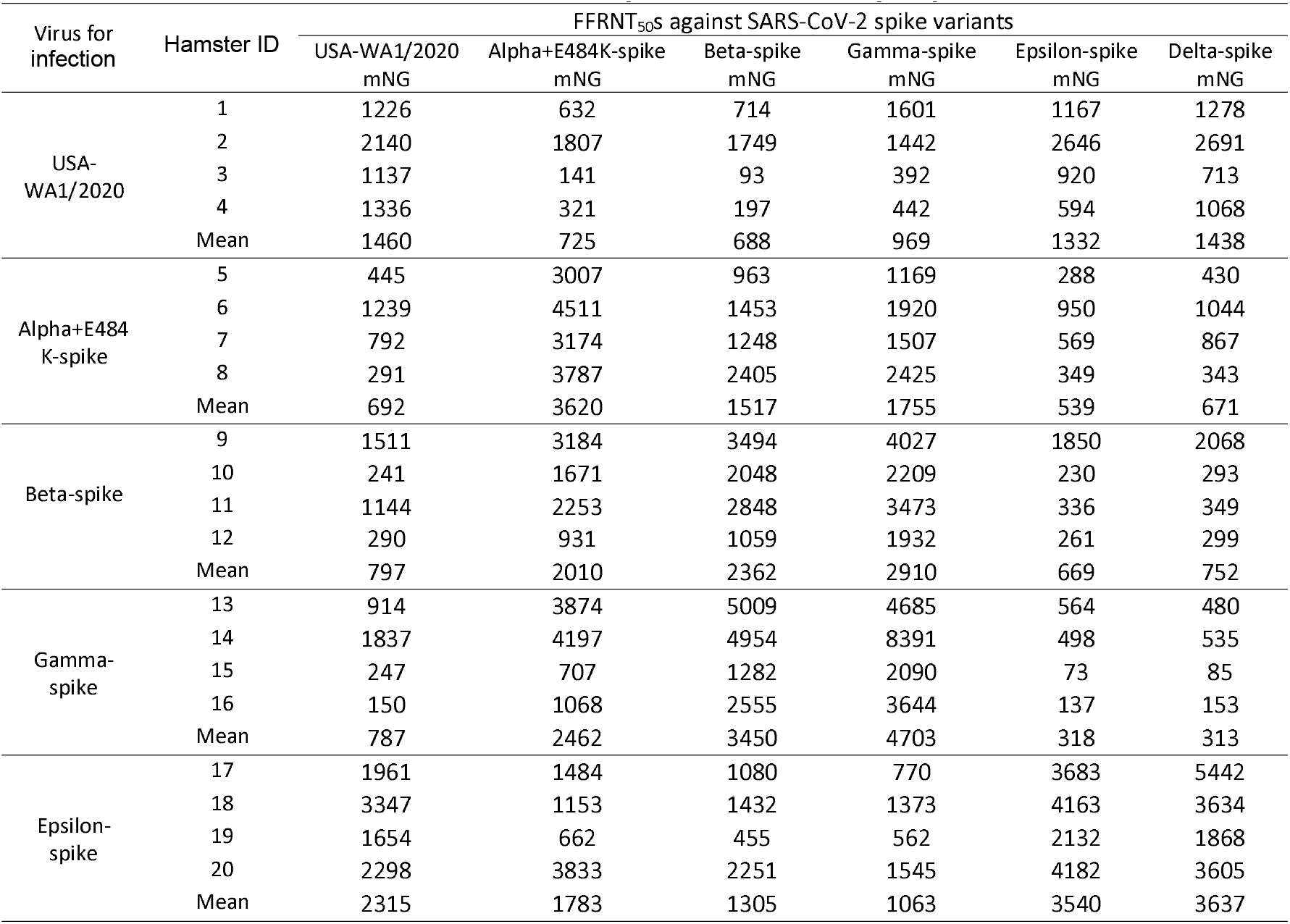
FFRNT_50_s of twenty hamster sera on day 28 post-immunization.

| Virus for infection | Hamster ID | FFRNT<sub>50</sub>s against SARS-CoV-2 spike variants |  |  |  |  |  |
| --- | --- | --- | --- | --- | --- | --- | --- |
|  |  | USA-WA1/2020 | Alpha+E484K-spike | Beta-spike | Gamma-spike | Epsilon-spike | Delta-spike |
|  |  | mNG | mNG | mNG | mNG | mNG | mNG |
| USA-WA1/2020 | 1 | 1226 | 632 | 714 | 1601 | 1167 | 1278 |
|  | 2 | 2140 | 1807 | 1749 | 1442 | 2646 | 2691 |
|  | 3 | 1137 | 141 | 93 | 392 | 920 | 713 |
|  | 4 | 1336 | 321 | 197 | 442 | 594 | 1068 |
|  | Mean | 1460 | 725 | 688 | 969 | 1332 | 1438 |
| Alpha+E484K-spike | 5 | 445 | 3007 | 963 | 1169 | 288 | 430 |
|  | 6 | 1239 | 4511 | 1453 | 1920 | 950 | 1044 |
|  | 7 | 792 | 3174 | 1248 | 1507 | 569 | 867 |
|  | 8 | 291 | 3787 | 2405 | 2425 | 349 | 343 |
|  | Mean | 692 | 3620 | 1517 | 1755 | 539 | 671 |
| Beta-spike | 9 | 1511 | 3184 | 3494 | 4027 | 1850 | 2068 |
|  | 10 | 241 | 1671 | 2048 | 2209 | 230 | 293 |
|  | 11 | 1144 | 2253 | 2848 | 3473 | 336 | 349 |
|  | 12 | 290 | 931 | 1059 | 1932 | 261 | 299 |
|  | Mean | 797 | 2010 | 2362 | 2910 | 669 | 752 |
| Gamma-spike | 13 | 914 | 3874 | 5009 | 4685 | 564 | 480 |
|  | 14 | 1837 | 4197 | 4954 | 8391 | 498 | 535 |
|  | 15 | 247 | 707 | 1282 | 2090 | 73 | 85 |
|  | 16 | 150 | 1068 | 2555 | 3644 | 137 | 153 |
|  | Mean | 787 | 2462 | 3450 | 4703 | 318 | 313 |
| Epsilon-spike | 17 | 1961 | 1484 | 1080 | 770 | 3683 | 5442 |
|  | 18 | 3347 | 1153 | 1432 | 1373 | 4163 | 3634 |
|  | 19 | 1654 | 662 | 455 | 562 | 2132 | 1868 |
|  | 20 | 2298 | 3833 | 2251 | 1545 | 4182 | 3605 |
|  | Mean | 2315 | 1783 | 1305 | 1063 | 3540 | 3637 |

**Extended Data Table 3.**
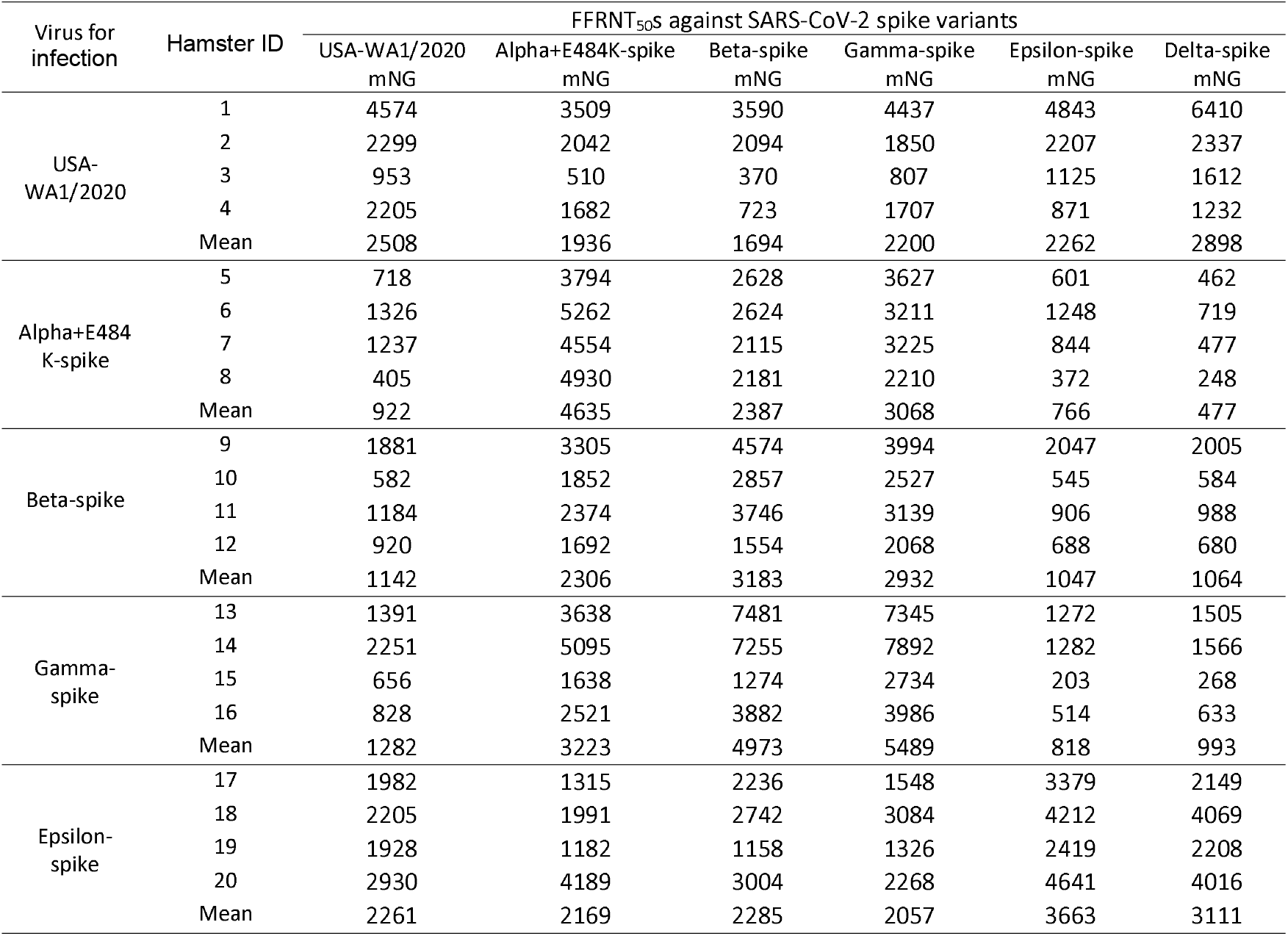
FFRNT_50_s of twenty hamster sera on day 45 post-immunization.

| Virus for infection | Hamster ID | FFRNT<sub>50</sub>s against SARS-CoV-2 spike variants |  |  |  |  |  |
| --- | --- | --- | --- | --- | --- | --- | --- |
|  |  | USA-WA1/2020 | Alpha+E484K-spike | Beta-spike | Gamma-spike | Epsilon-spike | Delta-spike |
|  |  | mNG | mNG | mNG | mNG | mNG | mNG |
| USA-WA1/2020 | 1 | 4574 | 3509 | 3590 | 4437 | 4843 | 6410 |
|  | 2 | 2299 | 2042 | 2094 | 1850 | 2207 | 2337 |
|  | 3 | 953 | 510 | 370 | 807 | 1125 | 1612 |
|  | 4 | 2205 | 1682 | 723 | 1707 | 871 | 1232 |
|  | Mean | 2508 | 1936 | 1694 | 2200 | 2262 | 2898 |
| Alpha+E484K-spike | 5 | 718 | 3794 | 2628 | 3627 | 601 | 462 |
|  | 6 | 1326 | 5262 | 2624 | 3211 | 1248 | 719 |
|  | 7 | 1237 | 4554 | 2115 | 3225 | 844 | 477 |
|  | 8 | 405 | 4930 | 2181 | 2210 | 372 | 248 |
|  | Mean | 922 | 4635 | 2387 | 3068 | 766 | 477 |
| Beta-spike | 9 | 1881 | 3305 | 4574 | 3994 | 2047 | 2005 |
|  | 10 | 582 | 1852 | 2857 | 2527 | 545 | 584 |
|  | 11 | 1184 | 2374 | 3746 | 3139 | 906 | 988 |
|  | 12 | 920 | 1692 | 1554 | 2068 | 688 | 680 |
|  | Mean | 1142 | 2306 | 3183 | 2932 | 1047 | 1064 |
| Gamma-spike | 13 | 1391 | 3638 | 7481 | 7345 | 1272 | 1505 |
|  | 14 | 2251 | 5095 | 7255 | 7892 | 1282 | 1566 |
|  | 15 | 656 | 1638 | 1274 | 2734 | 203 | 268 |
|  | 16 | 828 | 2521 | 3882 | 3986 | 514 | 633 |
|  | Mean | 1282 | 3223 | 4973 | 5489 | 818 | 993 |
| Epsilon-spike | 17 | 1982 | 1315 | 2236 | 1548 | 3379 | 2149 |
|  | 18 | 2205 | 1991 | 2742 | 3084 | 4212 | 4069 |
|  | 19 | 1928 | 1182 | 1158 | 1326 | 2419 | 2208 |
|  | 20 | 2930 | 4189 | 3004 | 2268 | 4641 | 4016 |
|  | Mean | 2261 | 2169 | 2285 | 2057 | 3663 | 3111 |

